# Insect Size Matters: Using Image and Dimensions Together Improves Image Classification

**DOI:** 10.1101/2025.06.27.661916

**Authors:** Melika Baghooee, Quentin Geissmann

## Abstract

Although visual features have been the cornerstone of insect recognition, morphological traits are often overlooked by modern “scale-invariant” deep learning-based methods. One such trait is absolute body size, which is commonly used by entomologists to describe species. We propose a novel approach that integrates explicit size information into computer vision models. Our method improves predictions of a Convolutional Neural Network (ResNet) by incorporating the likelihood of an insect belonging to each class based on its size and the size distribution of the training data. We compare the performance of the proposed size-aware approach against a standard ResNet and a feature-fused ResNet model, demonstrating that incorporating explicit dimensional traits, such as body size, can enhance classification accuracy across both balanced and imbalanced dataset scenarios. Our results also show that incorporating size reduces classification error, especially in low-data scenarios and for hierarchically coarse, taxonomic misclassifications (e.g., errors at the family level).

Despite being readily available and computationally inexpensive, size has, to our knowledge, not been used in this context before. We show that explicitly including size is both computationally efficient and practical, as it requires no specialized sensors and can be derived from standard image segmentation. This makes our approach highly scalable for automated monitoring of insects with camera traps. By acknowledging the importance of size, our approach also contributes to more robust and interpretable ecological modeling. This work opens new avenues for biologically informed AI applications in entomology and beyond.

## 1. Introduction

Traditional insect identification methods rely on physical examination and expert knowledge and can be limited by the availability of trained specialists (Badirli et al., 2023; Van Klink et al., 2022). Recent advancements in computer vision have enabled researchers to capture high-quality images of insects in the field, often using traps, digitising extensive museum collections, and engaging citizen scientists who contribute images to platforms like iNaturalist (Høye et al., 2021; Van Klink et al., 2022). Additionally, deep learning models, particularly convolutional neural networks (CNNs), have made automated insect identification more feasible by improving feature extraction, model generalization, and predictive accuracy (Van Klink et al., 2024). However, most of the automatic species identification tools rely solely on image data (Weinstein, 2018) whereas experts typically also use multiple contextual information (e.g., size, habitat, time of the year, etc.) to identify specimens more accurately.

Based on our review of some insect identification keys (e.g., Chinery & Gibbons, (2012)), we observed that overall insect size has long been used as an informative morphological trait to help entomologists differentiate visually similar species. In contrast, most models used in computational entomology are designed to be “scale-invariant” to ensure that the model can effectively analyze images at different scales. This often requires resizing the images to a fixed input scale to help maintain consistency in the features extracted across different scales (He et al., 2015a, 2015b; Simonyan & Zisserman, 2015). In this scaling, insects that are larger than the expected size should be downscaled, and the smaller insects need to be upscaled. As a result, CNNs receive only the resized inputs and have no access to the original size of the insect.

Table 1 shows the availability and use of explicit insect size in a selection of recent image acquisition and classification approaches. The methods were selected to represent some of the most recent techniques for each of the four types of emerging data outlined by Van Klink et al. (2022). For images collected by citizen scientists, like iNaturalist, as the pixel size vary between and within images, size cannot be trivially extracted and used for model training. However, looking at the standard imaging setups, such as smart camera traps and laboratory and museum setups, size information is often readily available as the camera-to-specimen distance is consistent. In these types of data, size can be approximated from images, yet, to our knowledge, only the study by Geissmann et al., (2022) has attempted to use size as additional input of the fully connected layer for classification. This highlights that, while size is readily available information for model training that could be trivially extracted from the images of such datasets, the effect of using size as additional information has not been fully explored.

**Table 1.**
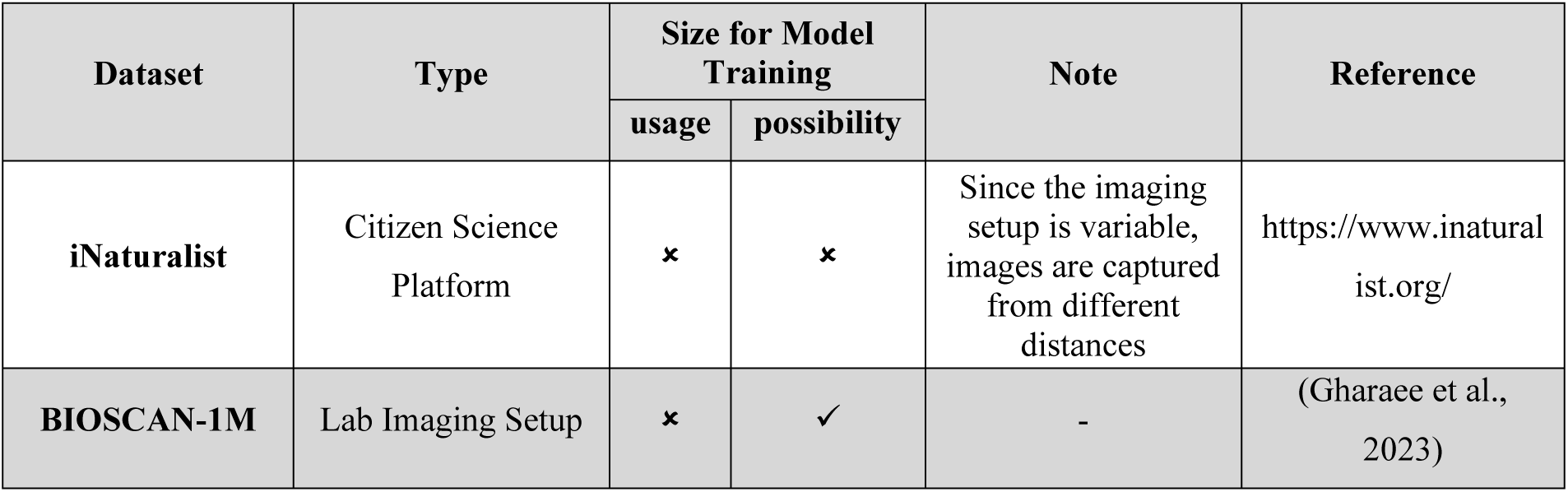
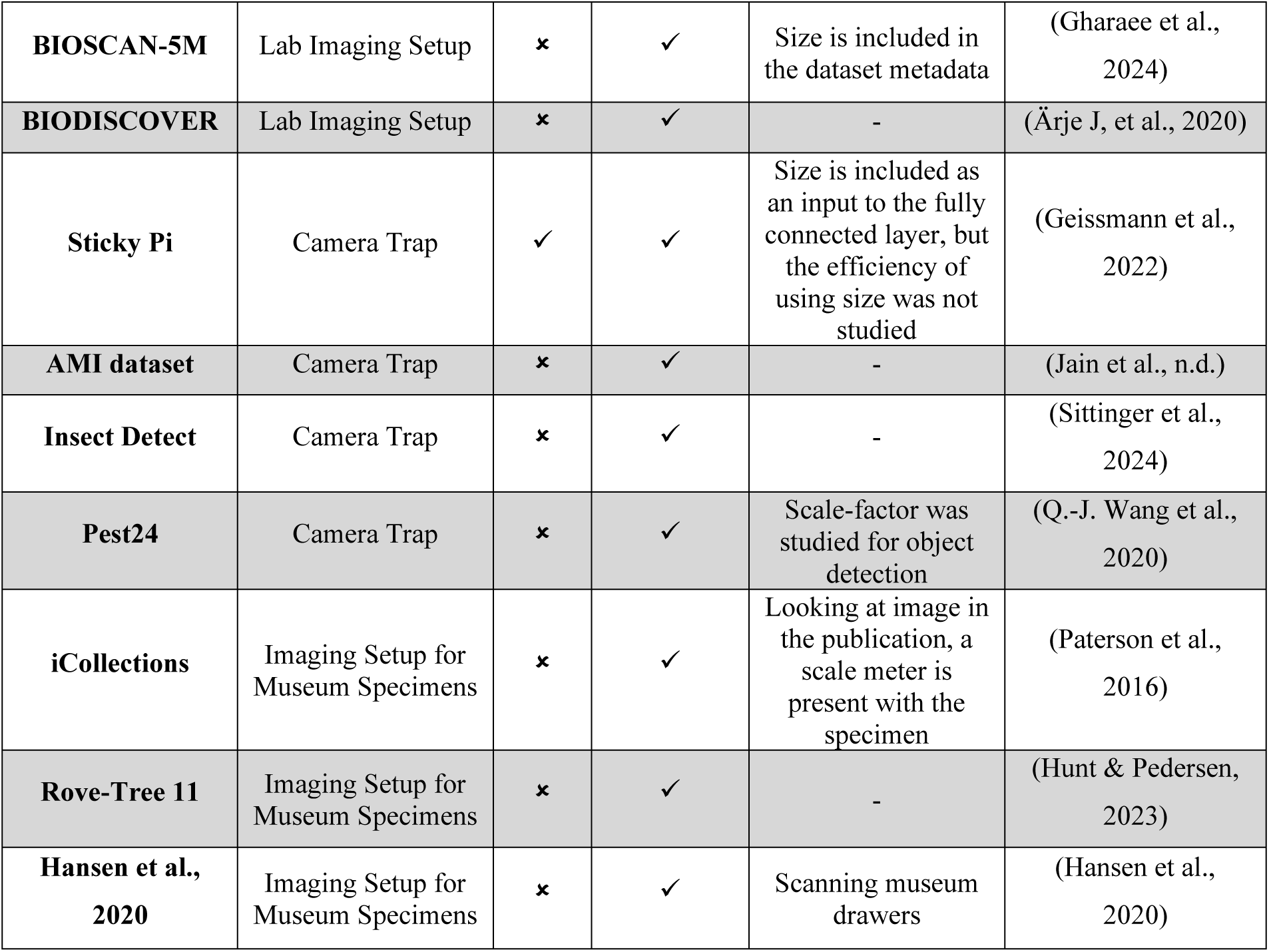
Assessment of 10 recent datasets, each using different imaging methods, to evaluate the availability of insect size information in the images, and whether this size data has been used as a feature in classification tasks

Another challenge with deep-learning models is their lack of interpretability, often referred to as the “black box” problem (Azodi et al., 2020). In contrast, additional features, such as size, observation time, and location, allow human experts to classify insects with greater confidence. Including these contextual details may also enhance deep learning models, potentially making them both more accurate and interpretable. Terry et al. (2020) have shown that incorporating metadata with computer vision models improves reliability in identifying insect species. Additionally, Gadzicki et al. (2020) studied the effect of the fusion of information from several data sources for the task of human activity recognition and found that fusion will most often improve performance by resolving ambiguity and improving the predictions provided from other reliable sources.

There are several strategies for multimodal fusion. Terry et al. (2020) proposed two primary approaches for incorporating metadata into computer vision models. First, in a combined approach similar to (Minetto et al., 2019) the metadata was concatenated with the extracted features from images before being fed into the fully connected layer. Second, with an ensemble method that treats the outputs of separate classifiers (one trained on image data and another on metadata) as relative probabilities. These probabilities are then combined (by product) to make final predictions. To our current knowledge, dimensional traits such as size are rarely incorporated in deep learning models for insect classification. Therefore, we suggest incorporating visual features along with explicit dimensional traits, such as size, whenever possible, in insect classification tasks. This approach can help enhance the accuracy of insect identification and provide a more detailed understanding of the model’s classification.

In this research, we hypothesize that a size-aware model will require fewer data to achieve the same level of performance as the standard one and is less affected by class imbalance since size is often an unambiguous and prominent variable. To test this hypothesis, we focus on the two mentioned fusion strategies: concatenating size information with visual features and combining predicted class probabilities with size-based likelihood via multiplication.

## 2. Methods

### 2.1 Dataset

In this study, we use the BIOSCAN-1M dataset (Gharaee et al., 2023). BIOSCAN-1M is a large collection of labeled insect images that have been taxonomically classified by experts. This dataset was selected for two key characteristics: (1) its standardized imaging system, which maintains a fixed camera-to-specimen distance across all specimens, and (2) the availability of a large and diverse sample of insects. These features make BIOSCAN-1M particularly suitable for insect size study. Among all the labeled species in this dataset, we initially filtered those with more than 130 available images. After this step, 85 species met the criteria. As this study focuses on insect size, we only used species with images of insects at the adult stage. Additionally, we manually curated this subset by removing images that featured: multiple insects in the image, no visible insects, or misclassified instances. Following the data cleaning process, 65 species remained for analysis. A description of the dataset is provided in Figure 1a-b, and examples of removed instances, together with the full taxonomic tree of the selected species, are presented in the Supplementary Materials. The genus *Megaselia* is presented as an example (Fig. 1c), illustrating the size distribution of species that are visually similar within a genus but differ in size.

**Figure 1.**
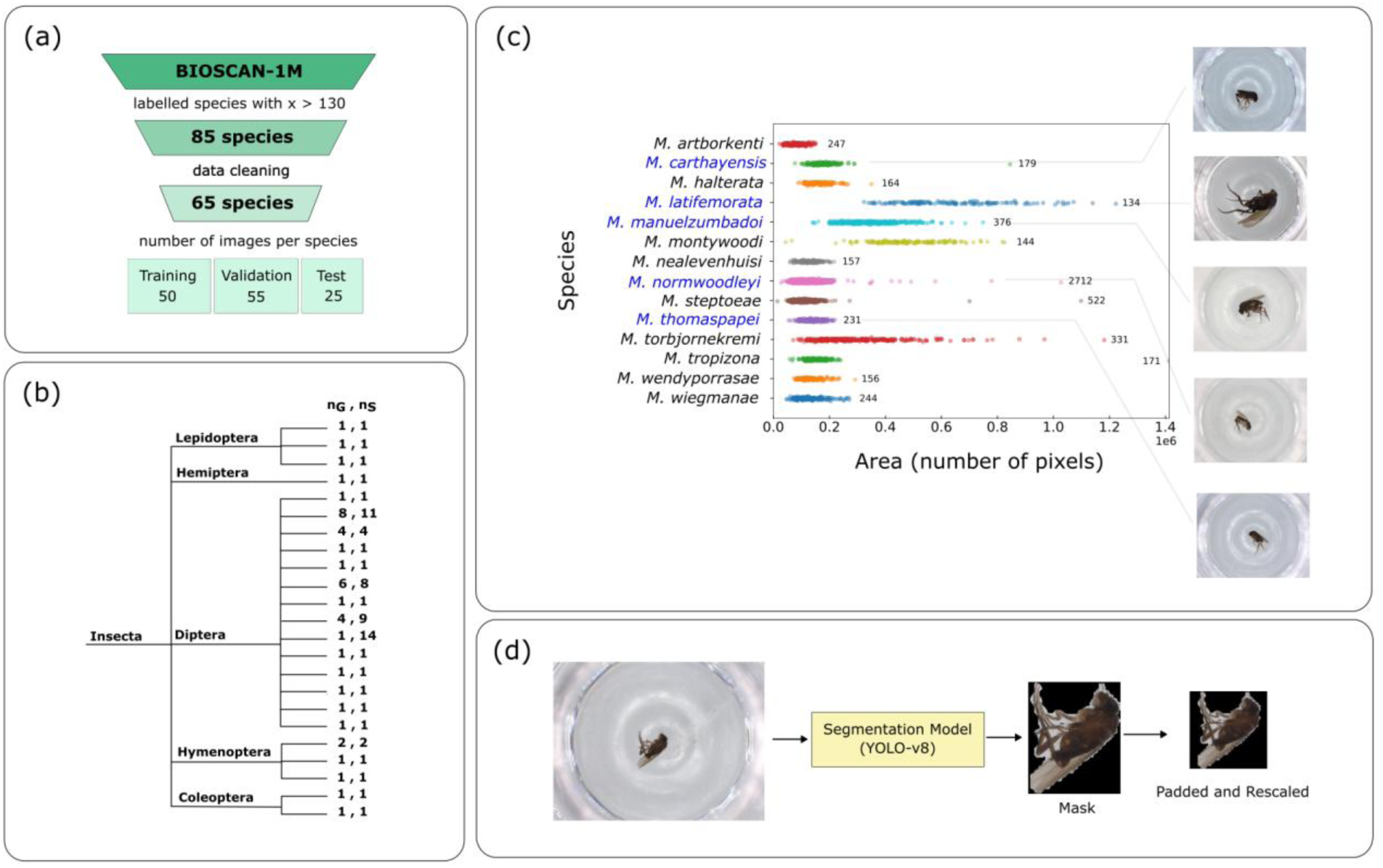
shows (a) data filtration and split, (b) the taxonomic tree of the selected species where 𝑛_𝑔_ = number of genera, and 𝑛_𝑠_ = number of species in each family. (c) shows, as an example, the distribution of the size of the insects within the genus *Megaselia*. The numbers written on the jitters show the number of available images for each species. The species written in blue are represented with an image showing the difference in size of visually similar species. (d) segmentation and data preprocessing

#### 2.1.1 Data Preprocessing

First, to prepare the images for the CNN model, each insect was segmented from its background (Fig. 1d). This was achieved by annotating 500 images, which were used to train a YOLO-v8 (Jocher et al., 2023) segmentation algorithm. The model validation showed high performance, with precision and recall values both stabilizing above 0.98. Using this model, new images were generated by applying the segmentation masks (setting the background to zero). The insect’s area was then defined as the number of non-zero pixels. For consistency in the rest of this manuscript, we will refer to size as the number of pixels in the insect’s mask.

#### 2.1.2 Experiments

After processing the images, they were split into training, validation, and test sets. Out of the 130 images available for each class, 50 were used for the training set, 55 for the validation set, and 25 for the test set. In this study, two types of experiments were investigated: (1) a balanced dataset and (2) an imbalanced dataset.

For the balanced dataset experiment, images were randomly sampled from both the training and validation sets, creating subsets with 5 to 50 images per species to explore how the size of the dataset affects the model’s performance, depending on the amount of data available for training.

In the imbalanced dataset experiment, we created random datasets with different levels of Shannon’s entropy (Shannon, 1948). Shannon’s entropy is calculated as follows:

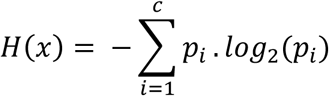

where:
𝑝*_i_*: the probability of each element *i* in the distribution given by 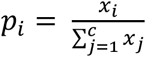 and 𝑥*_i_* represents the number of images in class *i* with 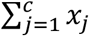 being the total number of images across all classes c: total number of classes

Each of the imbalanced cases was simulated using an exponential propensity-based sampling. Specifically, we considered a fixed number of total images of 1000, with a minimum of 5 and a maximum of 50 samples available per species. A set of imbalance severity levels using a parameter λ. For each λ, class-wise sampling probabilities were generated using the exponential decay function:

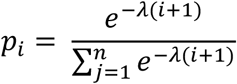

where:

𝑝_𝑖_: the sampling probability for class *i*

n: number of classes (65)

These probabilities were randomly shuffled to eliminate positional bias and used to sample a total of 1000 images across species. For each species, an exponential score was sampled for each image based on the assigned propensity. The top samples across all species based on this score were selected while ensuring that at least 5 images were present for each species. This introduced a controlled yet randomized species distribution for each imbalanced experiment.

A high entropy scenario occurs when the number of images per class is more balanced (for example, the highest entropy is when there are 5 images in all classes). A low entropy scenario, on the other hand, happens when few classes have a high frequency while others are rare (i.e., imbalance).

For both types of experiments, the test set was the same to have a fair and direct comparison.

### 2.2 Models

#### 2.2.1 ResNet

ResNet, short for Residual Network (He et al., 2015a) is an effective architecture for various computer vision tasks, including image classification, object detection, and segmentation, due to its unique use of residual blocks and skip connections. This design enables the model to learn deeper and more complex features without suffering from vanishing gradients. ResNet has also gained significant popularity in specialized domains, such as insect classification (Popescu et al., 2023). ResNet models require inputs of a fixed size (e.g., 224 × 224 pixels). This ensures that each image is treated consistently and is computationally consistent. For this reason, a ResNet-18 model is used as the baseline of a deep learning model, which is performant at classification, yet not “size-aware”.

#### 2.2.2 Size-awareness

As outlined in the introduction, there are several ways to integrate additional information, such as size, into a CNN architecture. In this study, we compare two approaches: (a) feature fusion and (b) Bayesian correction (Figure 2) with a standard ResNet. In the feature-fusion ResNet, size information is first normalized and then concatenated with the visual features extracted from the image, before being passed to the fully connected layer for classification. In our Bayesian Corrected (BC) ResNet approach, we use two components to make the final class prediction: (1) a ResNet-18 model that predicts based on the image alone, and (2) a size distribution model (similar to a Naive Bayes Classifier) for each class that was derived from the size of the insects in the training set. The size distribution model enables us to calculate the likelihood of each class for a given instance during inference. By assuming that the size of the insects in each class follows a Student’s t-distribution, the (log-)likelihood of an instance belonging to a class is calculated as follows (Jackman, 2009):

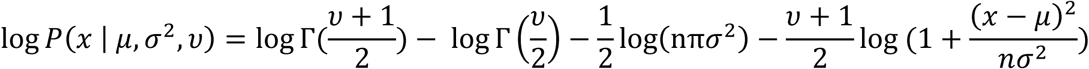

**Figure 2.**
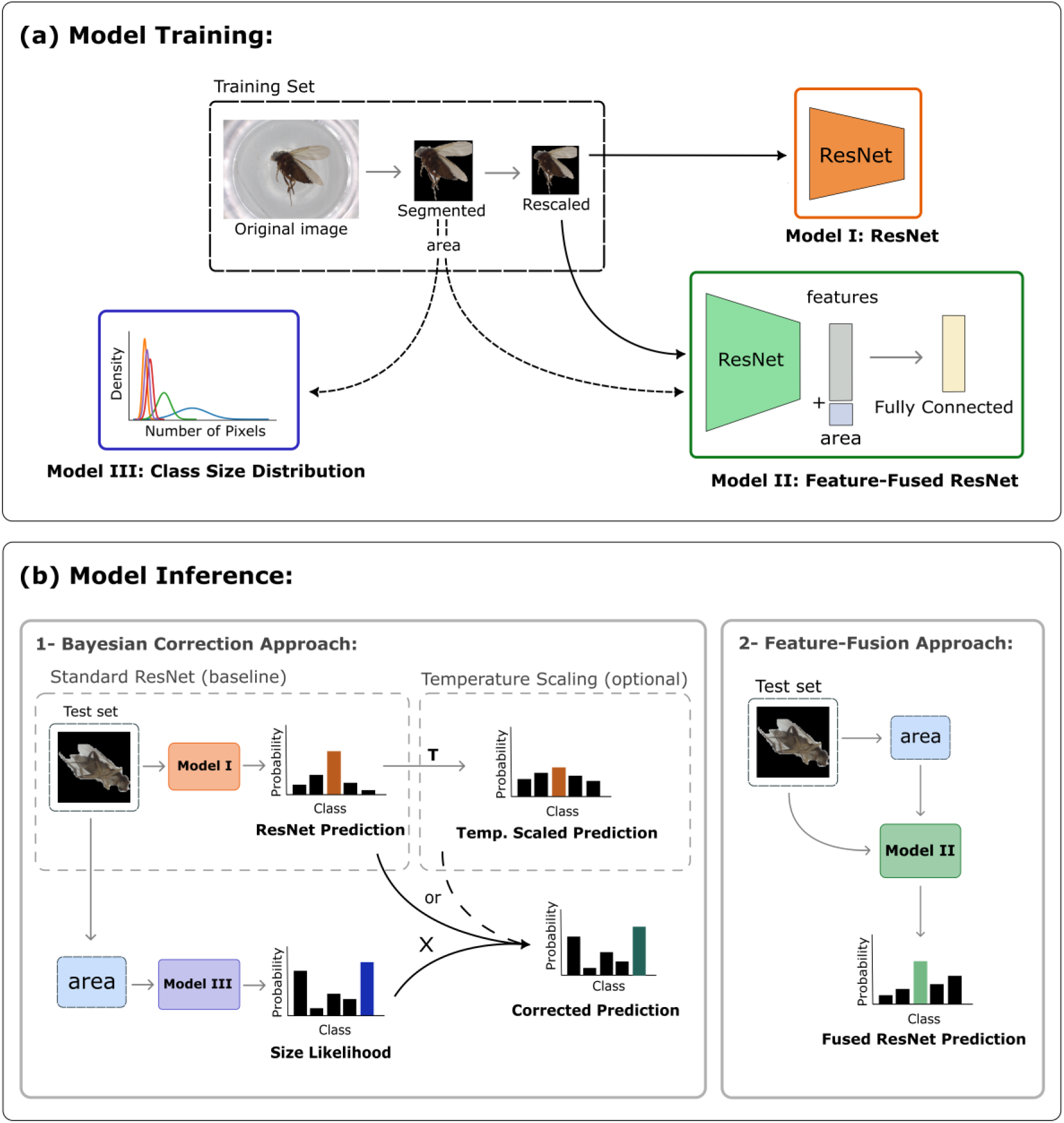
illustrates the Model Training and Model Inference phases: (a) the training phase, where original images are segmented and rescaled in preparation for ResNet training. The size (area) is obtained from the number of pixels belonging to the insect in the segmented masks and used both for the Fusion approach and to get the size distribution model. (b -1) the model inference phase for the Bayesian Correction approach, where the trained standard ResNet makes predictions on new data, and the predictions are either temperature scaled or directly adjusted based on the size likelihood obtained from the size distribution model (b -2) the inference of the Feature-Fusion approach

where:

x: the size of an instance (insect from the test set)

µ: the mean of the size of the insects in that class (approximated from the training set)

𝜎: the standard deviation of the size of the insects in that class (approximated from the training set)

n: number of training samples in that class

𝜈: degrees of freedom (𝑛 − 1)

In our approach, the final prediction is obtained by multiplying the ResNet model’s output with the prediction from the size distribution model. This ensures that size was effectively integrated into the final prediction of the model.

We assume the insects’ size to be normally distributed, while in nature, it might not be the case. To test this assumption, the size distribution for each selected species was evaluated using the Shapiro test. With a 0.05 significance level, only the size distribution of 10 out of 65 species was not normal, suggesting that the assumption holds reasonably well for most cases.

#### 2.2.3 Model Training and Evaluation

As illustrated in Figure 2, data from training and validation subsets based on the experiments explained in 2.2.1 were used to train two models: (1) a ResNet and (2) a size distribution model. During the training process, the best ResNet model, which had the lowest validation loss, was kept. To reduce overfitting, we used different types of augmentation, including random rotation values of (0, 90, 180, 270) along with horizontal and vertical flipping. A batch size of 16 was selected, and a step learning rate scheduler was used to gradually reduce the learning rate during training. The initial learning rate was set to 0.001, with a decay factor of 0.95, and the learning rate was updated after every epoch (step size = 1). The test set is the same among all experiments for a fair comparison. The size distribution model was developed based on the sizes of the images in the training set. Additionally, model training and evaluation were implemented in PyTorch (Paszke et al., 2019) (version 2.3.0) with torchvision (version 0.18.0), running on an external NVIDIA GeForce RTX 3080 Ti GPU.

To calibrate the model’s confidence, temperature scaling was used. According to (Guo et al., 2017) temperature scaling is an effective and simple calibration, especially on vision tasks. In temperature scaling, the logit layer of a deep learning model is rescaled by parameter T (temperature). The validation set is used to calculate the best value of T by minimizing negative log likelihood loss function respecting T, conditioned by T > 0 (Mozafari et al., 2019). To ensure that the model’s predictions were calibrated, we also tested a temperature-calibrated ResNet with the Bayesian correction.

Accuracy was used as an evaluation metric to compare the performance of the models. Accuracy is defined as the proportion of correct predictions across all instances. For the imbalanced dataset experiment, where class distribution varied in terms of the number of images per class, both macro and micro accuracies were computed to provide a more comprehensive evaluation.

Macro accuracy is calculated by averaging the class-wise accuracies. This metric accounts for an imbalance in class distribution, as each class contributes equally to the final accuracy score (independent of the number of images per class). On the other hand, micro accuracy computes overall accuracy based on the global counts of correct and incorrect predictions. This metric is sensitive to the size of the classes since it places more emphasis on the larger classes.

## 3. Results

### 3.1 Balanced Dataset Experiment

First, we wanted to evaluate how the different approaches performed and how data scarcity affected their performance. Figure 3a shows the difference between the accuracy of the approaches and a standard ResNet plotted against the number of training images per class. The approaches compared include the Feature-Fused ResNet, the Bayesian-Corrected (BC) ResNet, and the BC ResNet with temperature scaling. The results show that the Bayesian size-awareness approach (BC ResNet and BC ResNet with temperature scaling), outperforms both the standard ResNet and the Feature-Fused ResNet, especially when the training data is limited. For example, the BC ResNet trained on just 5 images achieved a gain of nearly 12.5% accuracy compared to a standard ResNet. Interestingly, temperature scaling did not yield consistent improvements over the standard Bayesian Corrected model. This may be due to the selective calibration of only the ResNet predictions, without calibrating the size distribution model, which could bias the uncertainty handling. The Feature-Fused ResNet did not perform consistently better, either. This happened possibly due to poor integration of the size feature with the visual features extracted by ResNet, which may have introduced confusion rather than useful signal (see the Discussion section). The absolute values of the accuracies of both ResNet, as the baseline, and BC ResNet, as the best approach, are presented in Figure 3b.

**Figure 3.**
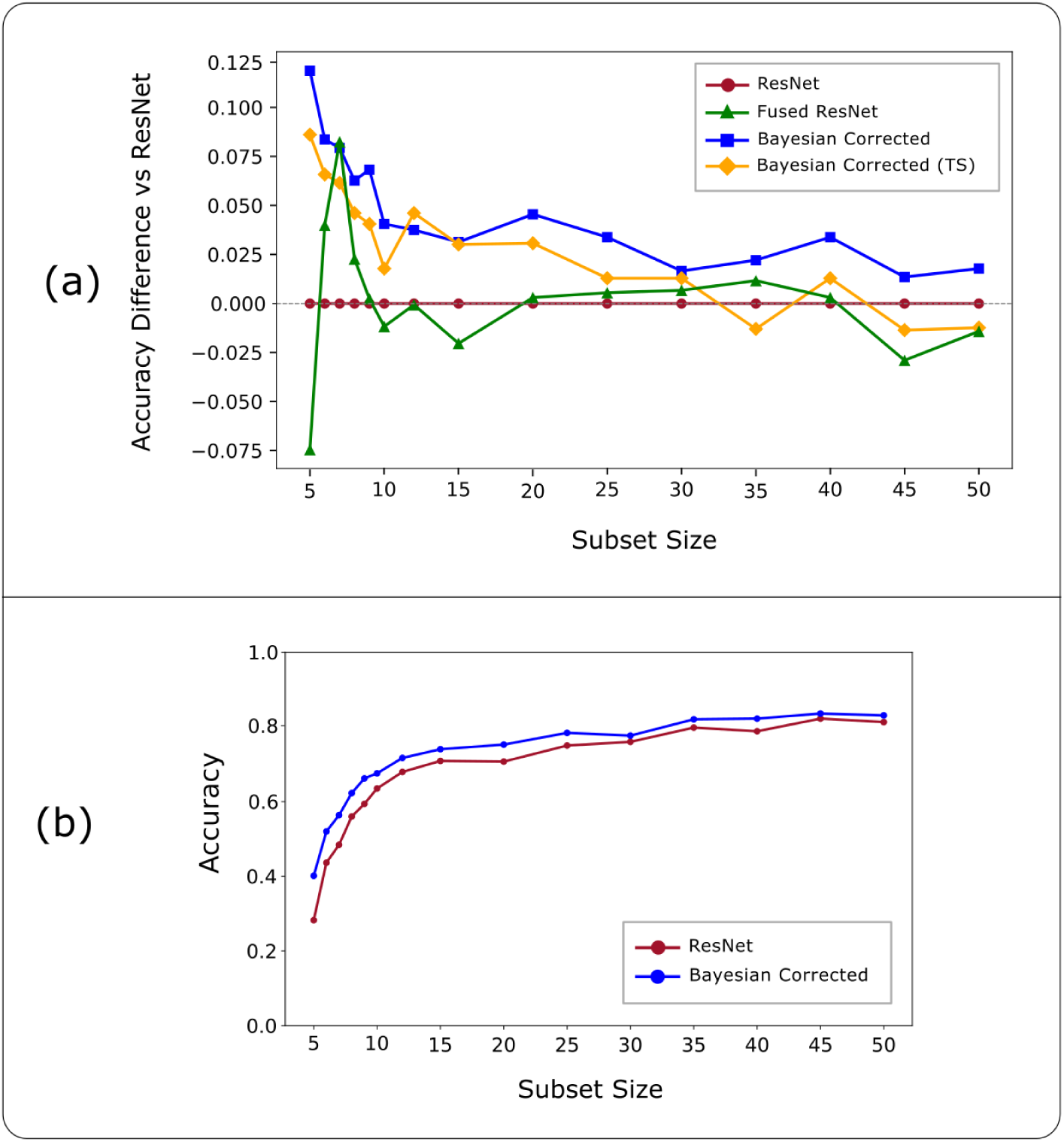
Performance of the models in the balanced dataset experiment, showing (a) the difference in accuracy of the standard ResNet, Fused ResNet, BC ResNet, and BC ResNet with temperature scaling across different numbers of training images per species (b) the absolute values of accuracy for the standard ResNet (baseline), and the BC ResNet (best model)

Overall, in the cases where fewer images were provided for the model to “learn” by the visual features, the size provides information beyond the image only. This is particularly valuable considering the challenges with classifying rare insect species, where it is difficult to obtain more data, but additional knowledge of size might be useful for the model to make a better prediction.

### 3.2 Imbalanced Dataset Experiment

Since the Feature-Fused ResNet failed to consistently outperform the standard ResNet, and the Temperature-Scaled Bayesian Corrected ResNet also did not consistently outperform the standard Bayesian Corrected ResNet (for reasons discussed in the Discussion section), we chose to proceed with the Bayesian Correction approach for the imbalanced experiments. To evaluate performance on the imbalanced dataset, both macro and micro accuracies were calculated, as these metrics provide different insights into model effectiveness. Figure 4 shows how the performance of each model is affected by changes in entropy. Looking at the slopes of micro accuracy (Fig. 4a), we see that as data imbalance increases, the standard ResNet experiences a slightly steeper decline in performance with a slope of -0.54 compared to the size-aware model (slope of -0.44). This suggests that the size-aware model is potentially better at handling class imbalance, likely because its prediction partially relies on the size information. However, statistical analysis indicates that this difference in slopes is not significant (p > 0.1). While a significant difference in intercepts (p < 0.01) confirms that the size-aware model performs better overall, by consistently achieving higher micro accuracy than the standard ResNet, even in the presence of class imbalance.

**Figure 4.**
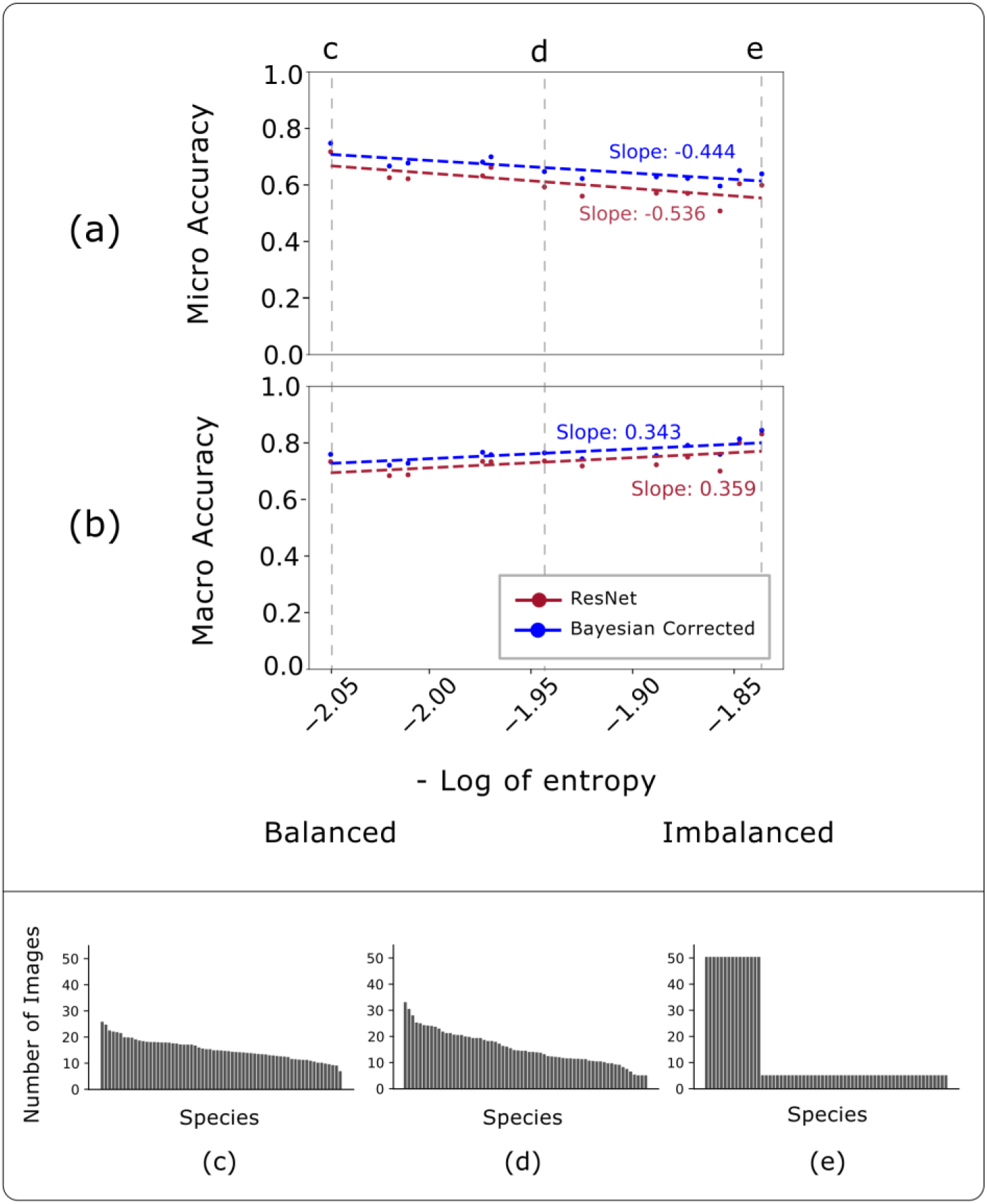
Performance of models on datasets with varying species balance: (a) Micro accuracy: Highlights the influence of dominant classes on overall prediction accuracy (b) Macro accuracy: Demonstrates that the BC model is less affected by class imbalance compared to the standard ResNet (c) High-balanced dataset: Distribution where the number of images per species falls within a narrow range (d) Medium-balanced dataset: Shows a broader range in image counts per species, indicating moderate imbalance (e) Highly imbalanced dataset: A few species dominate, contributing up to 10 times more images than others

Conversely, the macro accuracy metric, which aggregates correct predictions across all classes, does not fully capture the challenges posed by data imbalance. For the imbalanced cases, the model can learn to “cheat” by predicting the class with more data, more often, and looking at the overall correct predictions is not as insightful as considering class-wise accuracy. The results shown in Figure 4b also show this increase in macro accuracy of both models as the imbalance increases. Although the difference in slopes remains statistically non-significant, the size-aware model again has significantly higher intercepts (p < 0.05). Overall, the performance trends of both models are consistent, with the size-aware model outperforming the standard ResNet.

### 3.3 Effect of Size-Awareness on Taxonomic Error

To examine how using size distribution information impacts model performance at different taxonomic levels, we looked at the number of errors on each taxonomic level (Figure 5). By comparing models with and without size correction, this figure provides insight into where and how size information helps disambiguate classes. To get this information, we looked at the predicted species on both ResNet and BC ResNet and referred to the taxonomic information in the metadata of each of the images in the dataset. If the predicted species were within the same genus, the severity of the error is less, but if it is from a different order, it is a more severe error.

**Figure 5.**
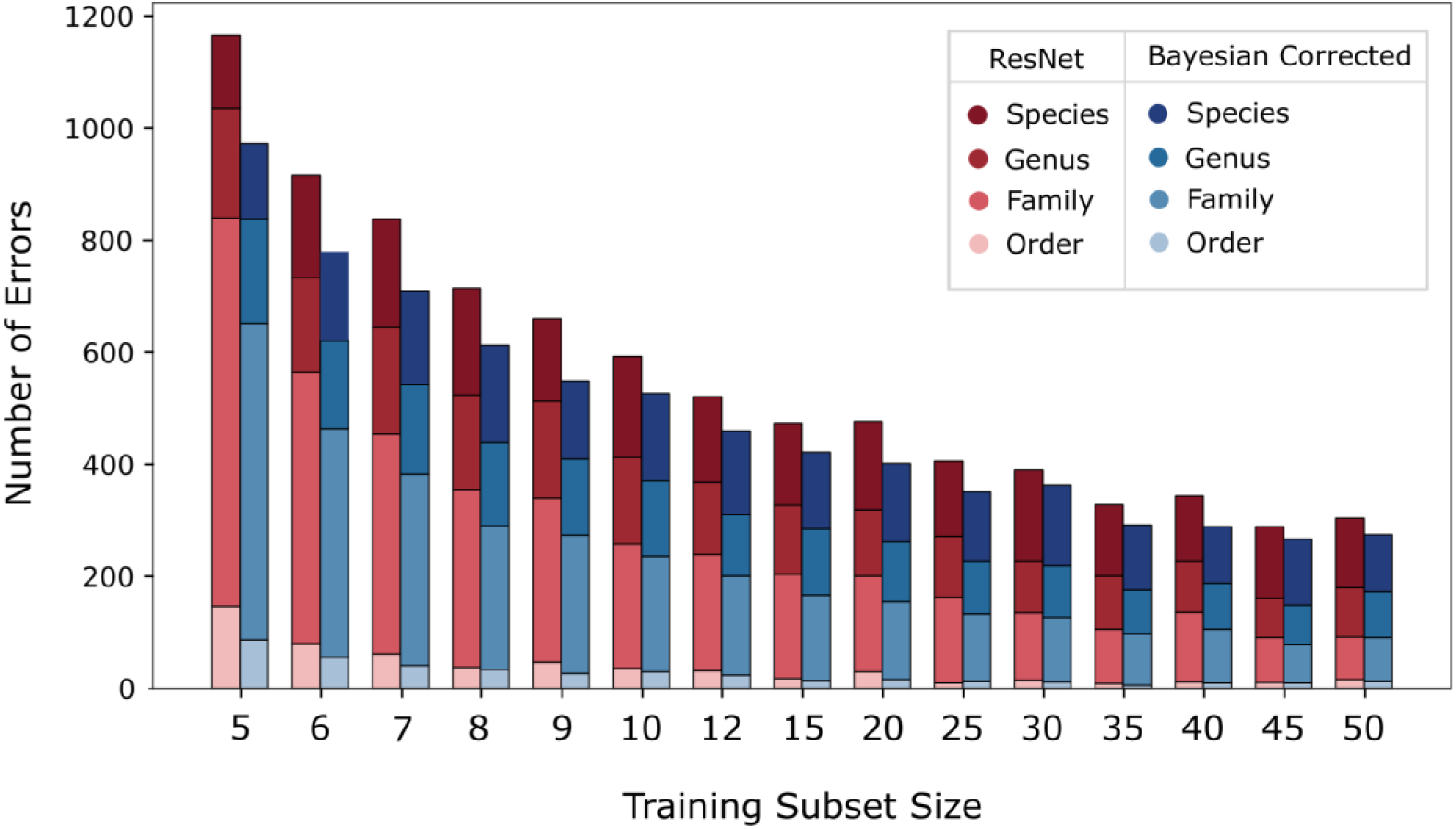
Illustrates the number of classification errors across different taxonomic levels (Order, Family, Genus, and Species) as a function of training subset size.

The BC ResNet consistently results in fewer errors across all training sizes. This means that by knowing the size, the model is making fewer errors, especially in cases where the number of images in the training set was not enough for the model to learn from the visual features of, e.g., five images. At coarser taxonomic levels like Order and Family, the BC ResNet shows more improvements, likely because size is a strong cue at higher taxonomic levels and for cases where insects are visually ambiguous, knowing the size helps reduce the errors.

At finer taxonomic levels (Genus and Species), the performance gap between the models is narrower. This indicates that, while size still provides a small regularizing benefit, visual features dominate decision-making at finer taxonomic levels. Nevertheless, even here, the BC ResNet tends to perform at least as well as, if not better than, the baseline.

These results highlight that incorporating size information primarily helps resolve ambiguity where taxonomic categories are visually less distinct, and training data is limited. They also show that the addition of size, as prior knowledge, can be especially beneficial at coarser levels and in low-data scenarios. It also justifies the use of size-aware corrections in applications where coarse classification is a priority, such as automated biodiversity monitoring or specimen sorting.

## 4. Discussion

This study shows how using explicit dimensional traits, such as size, can enhance model performance. While size is often readily available and virtually free to use, its effectiveness when incorporated into a deep learning model has not been widely explored in this context. Our “Bayesian Corrected size-aware” approach can achieve acceptable performance with little data and is less affected by class imbalance. These two address two major challenges with training deep learning models for insect classification: data hunger and imbalance. Importantly, our approach also offers greater flexibility than black-box late fusion methods, as it enables both training and inference even when size information is unavailable for some images. Moreover, it can be applied to already trained models without requiring retraining. This section discusses the implications of these findings, potential applications in biodiversity research, challenges encountered, and directions for future research.

### 4.1 Value of size awareness in limited-data scenarios

The results from the balanced dataset experiment underscore the size-awareness advantages when training data is limited. The Bayesian-Corrected approach can generalize better with limited samples, which is particularly useful in applications like rare species classification when gathering large datasets is challenging. The size distribution of classes requires two main information, the class’s mean and standard deviation. These parameters can be derived with relatively few data. Of course, more data helps approximate more accurate distributions, but it is still easily obtained even with fewer data (e.g., 5 individuals in this study). By using the size distribution model as additional knowledge, the models have an additional predictive feature, which can compensate for data scarcity. A key advantage of this approach is that it remains flexible when size data is unavailable (such as in mixed or incomplete datasets), as it can be seamlessly ignored without altering the model structure compared to a feature-fused approach. It can also extend to other additional, easily obtained information (e.g., in case of images obtained from camera traps, the time of the day and the season the image was captured), or even external textual descriptions and trait databases, all of which could potentially enhance model performance.

### 4.2 Robustness to class imbalance

The imbalanced dataset experiment shows the robustness of the size-aware approach to class distribution. While both models experience performance degradation with increasing imbalance, the size-aware approach maintains consistently higher accuracy levels. This robustness is essential in ecological datasets, which often contain a mix of common and rare species. It indicates that the Bayesian correction with size could help better species identification in these contexts. This robustness is achieved because of the availability of size-based priors, which provide a more stable form of contextual information, helping the model reduce reliance on implicit assumptions about class frequency.

### 4.3 Challenges and Limitations

While these findings are promising, there are still challenges associated with size-informed corrections. One key limitation is the applicability of this approach to datasets derived from citizen science platforms. Since different users take images from different distances, accurately defining insect size can be challenging. Despite this challenge, the size-correction method remains useful for classifying insects in datasets with standardized imaging setups, similar to the BIOSCAN (Gharaee et al., 2023), and arguably, for smart camera trap data, such as Sticky-Pi (Geissmann et al., 2022), and Insect Detect (Sittinger et al., 2024).

In this study, the insects’ size was assumed to be normally distributed, however it is important to keep in mind that not all insect classes in nature might be normally distributed. In such cases, a possible solution is to model size as a mixture of Gaussians, which can better capture more complex distributions. Additionally, defining size as the area (i.e., number of pixels) can be highly sensitive to the insect’s pose in the image, introducing greater variability and potentially resulting in long-tailed distributions that reduce the reliability of size-based likelihood estimates. A more accurate definition and estimate of the distribution of the size could help improve the model’s performance

From a biological perspective, while this approach can be very useful in cases with visual similarity and significant size difference, there are cases where modeling size with a simple Gaussian distribution may not be sufficient to model size accurately. For insects that the size is different between males and females (sexual dimorphism) (Stillwell et al., 2010) using the size can be a challenge. Additionally, in seasonal polyphenism, where insects undergo size changes in response to environmental conditions such as temperature and time of year. Insects also exhibit diet-mediated phenotypes, as seen in some caterpillar morphs and the castes of social insects (Simpson et al., 2011).

Despite these challenges, the concept of integrating other sources of information into an image-based deep-learning model remains promising. For example, (Terry et al., 2020) showed that incorporating additional metadata along with images improved the performance of their classification model. This suggests that combining multiple data sources could significantly improve the accuracy of image-based insect classification models. Furthermore, since size is often readily available and nearly free to obtain, it provides an excellent opportunity to enhance model performance without significant added cost or complexity.

### 4.4 Future directions

Future research could explore the addition of other traits, such as habitat information or seasonal occurrence, to address some of the challenges outlined earlier. These supplementary features may help further refine the accuracy of size-informed corrections and provide more robust classifications. Moreover, integrating these variables could provide valuable insights into the decision-making process of the model, changing the deep learning classification task from a “black box” into a “grey box,” where the model’s predictions become more interpretable, which is an ongoing research effort (Azodi et al., 2020; Hassija et al., 2024).

## Acknowledgements

The authors were both supported by the Novo Nordisk Foundation Start Package grant (NNF22OC0077040).

## Author Contribution

M. Baghooee and Q. Geissmann conceived the ideas; M. Baghooee curated the data; M. Baghooee and Q. Geissmann contributed to the implementations; M. Baghooee and Q. Geissmann analyzed the results; M. Baghooee led the training and evaluation of the models; M. Baghooee led the data visualization; M. Baghooee led the writing; Q. Geissmann reviewed the first draft; Q. Geissmann was the project manager and supervisor. Both authors contributed critically to the drafts and gave final approval for publication.

