## supplementary-materials for "Insect Size Matters: Using Image and Dimensions Together Improves Image Classification"

### Supplementary Material:

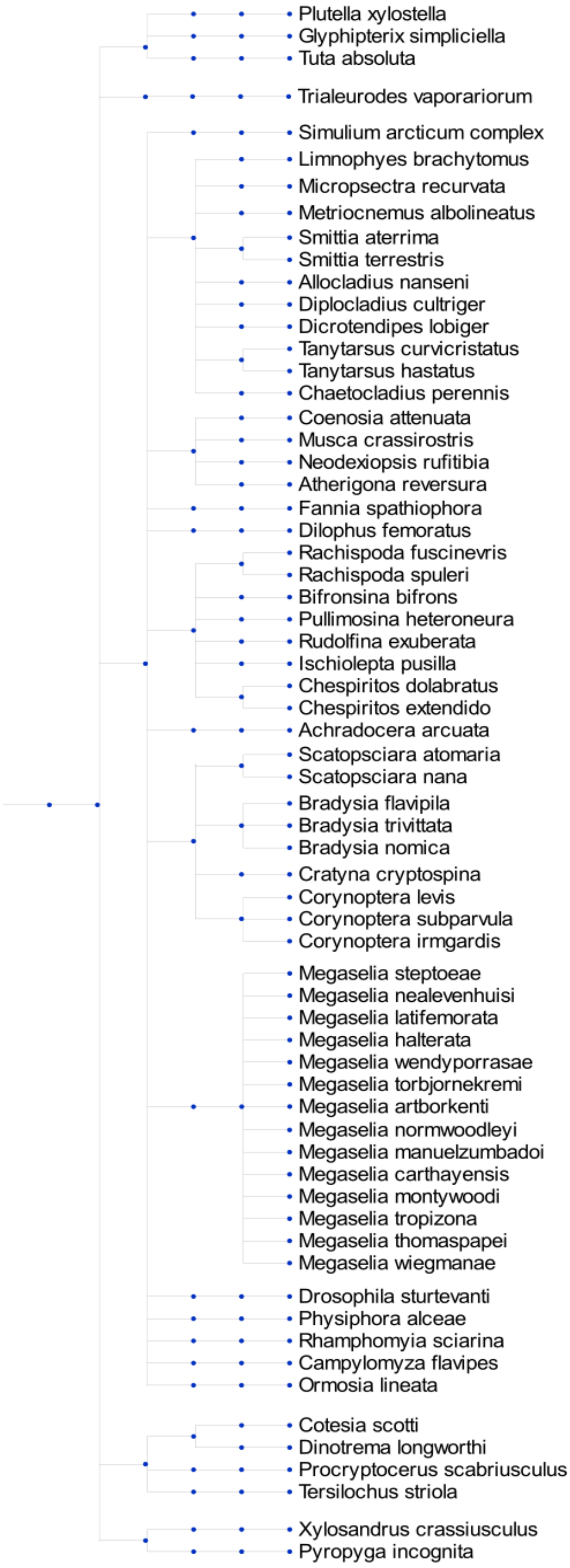


Fig. I- The taxonomic tree of the dataset used for this study

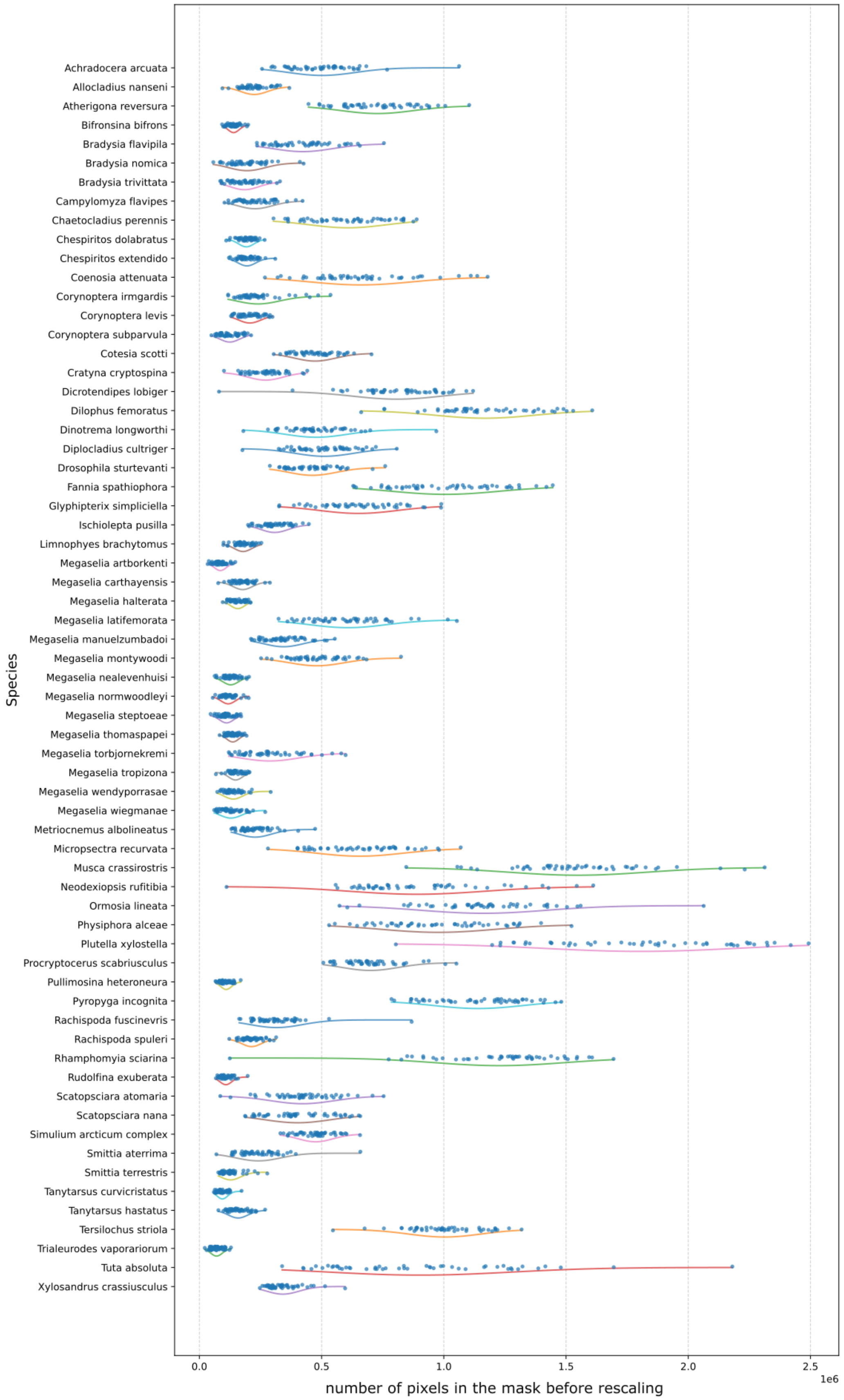


Fig. II- The distribution of the size (number of pixels before rescaling) of all species in this study

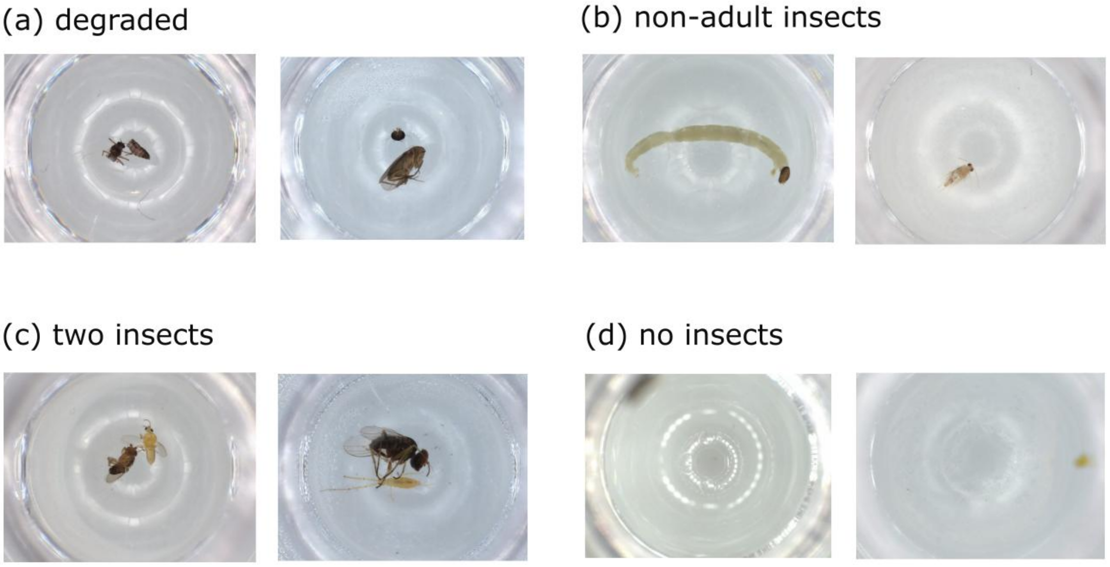


Fig. III- Examples of removed data during data cleaning

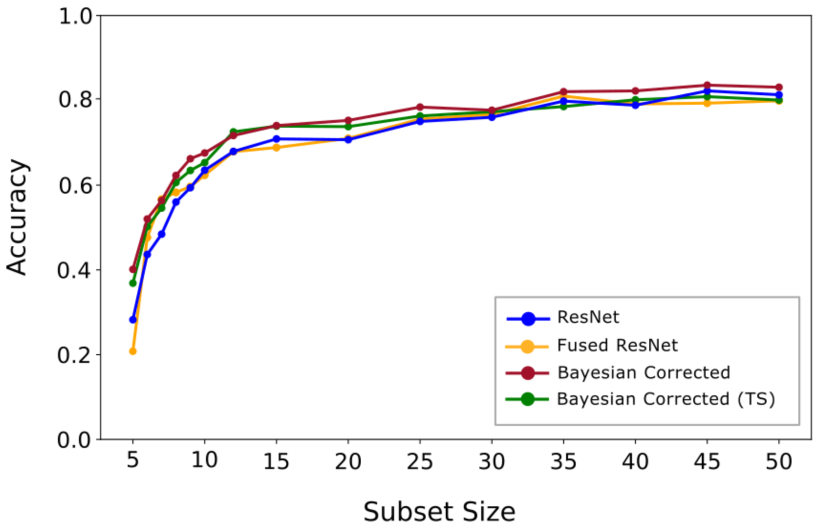


Fig. IV- Performance of all models in the balanced experiment
